# arakno - An R Package for Effective Spider Nomenclature, Distribution and Trait Data Retrieval from Online Resources

**DOI:** 10.1101/2021.09.30.462516

**Authors:** Pedro Cardoso, Stano Pekar

## Abstract

Online open databases are increasing in number, usefulness, and ease of use. There are currently two main global databases exclusive for spiders, the World Spider Catalogue (WSC) and the World Spider Trait (WST) database. Both are regularly used by thousands of researchers. Computational tools that allow effective processing of large data are now part of the workflow of any researcher and R is becoming a *de facto* standard for data manipulation, analysis, and presentation. Here we present an R package, *arakno*, that allows interface with the two databases. Implemented tools include checking species names against nomenclature of the WSC, obtaining and mapping data on distribution of species from both WST and the Global Biodiversity Information Facility (GBIF), and downloading trait data from the WST. A set of tools are also provided to prepare data for further statistical analysis.

---

Online open databases are increasing in number, usefulness, and ease of use. In this respect, spider researchers have always been at the forefront, with the World Spider Catalogue (2021; hereafter WSC) serving as a continuously updated reference on the nomenclature of the group. WSC provides taxonomic information including the most updated nomenclature, previous nomenclature, synonyms or misidentifications in past taxonomic works. The WSC also allows registered users to freely and immediately download any publication of taxonomic relevance, a privilege that almost no other researchers have. Including all spider species ever described, currently close to 50 000, the WSC is and will continue to be extensively used for multiple purposes within and outside research, from checking current nomenclature in faunistic works, to serving as the taxonomic basis to major conservation efforts worldwide (e.g. Seppala et al. 2018). Its data are free to use and already in 2015 logged a daily average of 600 hits and 400 downloads (Nentwig et al. 2015).

The World Spider Trait database (Pekár et al. 2021; hereafter WST) is a recent addition to the arsenal of virtual infrastructures available to us. It intends to compile curated data on spider traits (such as morphological, ecological, physiological, or behavioural) and this way both serve as a repository for data used in many publications and promote the use of trait data for eco-evolutionary analyses (Lowe et al. 2020). Its data are provided under a CC BY 4.0 license, and given its extremely recent availability no relevant statistics are available yet.

Both databases store huge amounts of data that can be quickly accessed via online applications (see https://wsc.nmbe.ch/ and https://spidertraits.sci.muni.cz/, respectively). These applications provide simple tools for data retrieval if the amount of data to be retrieved is relatively small. However, as the number of species of interest increases, as in, for example, community ecology studies, the use of online tools becomes ineffective.

Here we present the R package *arakno* (ARAchnid KNowledge Online; Cardoso 2021) available from CRAN (https://CRAN.R-project.org/package=arakno). It includes a suite of functions within the R environment (R Core Team 2021) allowing to download data from both WSC and WST, checking valid species names, listing or mapping species distributions, and preparing trait data for further analysis. The R language is becoming a *de facto* standard for data manipulation, analysis, and results presentation in, e.g., ecology and evolution, and we expect these functions will be of use for many arachnologists, among other researchers. Specifically, taxonomists or community ecologists can use them to assemble lists of valid species names with authorities and family membership or to produce maps of current known distribution. Evolutionary biologists, community ecologists, macroecologists, among many others, can download trait data to test predefined hypotheses.

The package *arakno* contains the following functions:

*wsc()* – Downloads the current, daily updated, list of valid spider names with their distribution data (list of countries) from WSC and returns a *data*.*frame* named *wscdata* to the global R environment.

*checknames(tax, full = FALSE, order = FALSE)* – Compares binomial species, genus and family names provided in a vector named *tax* with the nomenclature from WSC and, in case mismatches are detected, returns a four column array with original names (Species), possible matches (Best match; Alternative match(es)) and a Note indicating whether a nomenclature change, junior synonym or misspelling was detected.

Misspellings in names are matched using fuzzy logic (minimum Levenshtein edit distance, representing the minimum number of character deletions, additions or substitutions needed to convert one word into another). Subspecies names are not eligible. If *full* = *TRUE* all species names are returned, including valid names. As in other functions of this package, the results can be ordered alphabetically or follow the order given in *tax* (the default).

*authors(tax, order = FALSE)* – Returns authority and year for binomial species names provided in *tax* according to the WSC. If genera or families are used, data on all their species are retrieved.

*distribution(tax, order = FALSE)* – Returns distributions (countries or geographic areas) of species given in *tax* from the WSC. If genera or families are used, data on all their species are retrieved.

*lsid(tax, order = FALSE)* – Returns the *LSID* (Life Science Identifier, a unique code for each species) of species given in *tax* from the WSC. If genera or families are used, data on all their species are retrieved.

*species(tax, order = FALSE)* – Returns all species currently belonging to a family or genus given in *tax* according to the WSC.

*taxonomy(tax, check = FALSE, aut = FALSE, id = FALSE, order = FALSE)* – Returns the classification at sub/infraorder, family, and genus level for species in *tax* according to the WSC. If genera or families are used, data on all their species are retrieved.

Optionally, if *check = TRUE* the function will check the names (similar to function *checknames*); if *aut = TRUE* the function will provide authorities (similar to function *authors*); and if *id = TRUE* the function will give the LSIDs from the WSC (similar to function *lsid*).

*traits(tax, trait = NULL, sex = NULL, life = NULL, country = NULL, habitat = NULL, user = “”, key = “”, order = FALSE)* – Downloads all available trait data from the WST for all taxa given in *tax*. If genera or families are used, data on all their species are retrieved. Eligible values for *trait, sex, life, country*, and *habitat* are available at *https://spidertraits.sci.muni.cz/traits*. For traits, the abbreviation is used. To access embargoed data, a *user* name and API (Application Programming Interface) *key* from the WST must be provided.

*records(tax, order = FALSE)* – Returns coordinates and sources of occurrences for the taxa given in *tax* from both the WST and the Global Biodiversity Information Facility (GBIF; GBIF.org 2021). GBIF allows easy access to distribution data from multiple institutions worldwide, with the licensing being determined for each dataset. If genera or families are used, data on all their species are retrieved. The user should be aware that coordinate data returned are incomplete, often suffering from the Wallacean shortfall, i.e., incomplete geographical coverage (Lomolino 2004; Cardoso et al. 2011).

*map(tax, countries = TRUE, records = TRUE, hires = FALSE, zoom = FALSE, order = FALSE)* – Returns a map of distribution for the species given in *tax* using a list of *countries* and regions according to the WSC and/or *records* with coordinates available from the WST and GBIF. If genera or families are used, data on all their species are retrieved. Maps can be of high resolution (*hires = TRUE*) and the area zoomed (*zoom = TRUE*) to the area covered by the *records* available for each species.

Use of all functions is demonstrated in Appendix 1. Additional functions are planned to be implemented in future releases of the package *arakno*. Specifically:

1. Connection with other arachnid databases, such as the World Catalog of Opiliones (Kury et al. 2020). This will only be possible when digital identifiers and APIs are available, so we would like to appeal to authors of the catalogs to implement these requirements.
2. Connection with the International Union for the Conservation of Nature Red List (IUCN 2021) once arachnid coverage is comprehensive.
3. Seamless integration of *arakno* with other packages developed by the same author for ecology, biogeography and conservation purposes (Cardoso et al. 2015; Cardoso 2017), for an improved workflow.

The future development and utility of functions within the package are dependent on maximizing species and trait data coverage, data standardization and enable connectivity of databases according to the FAIR principles - Findability, Accessibility, Interoperability, and Reuse (Wilkinson et al. 2016). By following these principles, it is possible to easily connect *arakno* with multiple databases independently of their specific goal or format, this way maximizing data usage. We therefore strongly encourage:

1. Arachnid database managers to follow the FAIR principles so that data can be fully explored for multiple purposes.
2. Arachnid experts to contribute to the multiple databases (e.g. Pekár et al. 2021) to both guarantee archiving and re-use of data, and contribute to global efforts on species knowledge and conservation.

The R package *arakno* tries to fill a gap in facilitating access to online open databases using robust, flexible, and expandable functions. It is open source and any contributions and suggestions to its continuous development are most welcome.

## Acknowledgments

We thank all the developers and data providers of the WSC and WST. Adam Kučera for development of the WST API. The Pentti Tuomikoski Grant from the Finnish Museum of Natural History provided financial support to the development of *arakno* and the WST.

## Appendix 1

Below we show the utility of the functions from the ***arakno*** package in three examples that are mimicking common situations. The package can be loaded with the following command:

~~~
> **library(arakno)**
~~~

### Example 1

This example shows how to find valid species names, authorities, LSID codes and prepare this information in a table, which is a common requirement of papers on community ecology, for example.

You may store the list of names that you want to check in an Excel spreadsheet. Copy the column with names to R via clipboard and run the following command:

~~~
**> tax <-scan(“clipboard”, what=“char”, sep=“\t”)
> tax**
[1] “Mangora acalypha” “Clibiona compta” “Pardossa amendata” “Gyrakancium eraticum”
[5] “Theridion impressum” “Zodarum rubidum” “Agelena similis” “Arraneus”
~~~

The list contains eight names, the first is a valid name, the second is a valid name with a single typo, the third is a valid name with two typos, the fourth has multiple typos, the fifth is a species that changed genus, the sixth is a junior synonym with a typo, the seventh is a junior synonym of a species that changed genus, and the eight is a genus name with a typo. To check the validity of names use function `checknames` and place them into an object named e.g. `tax1`, as it will be used later:

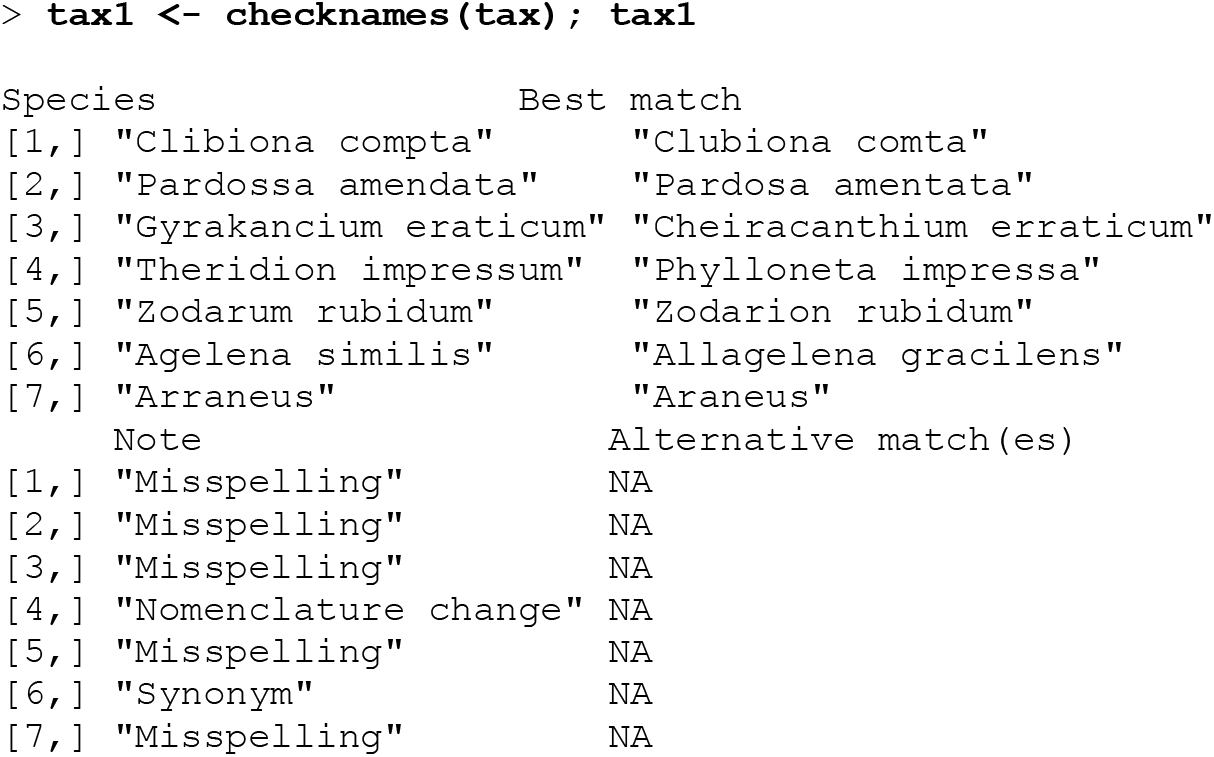

The function returned four columns, the first containing the list of original but incorrect species names (the first one was omitted as it is correct), the second one lists possible best matches, the third the reason for mismatch, and the fourth alternative match(es). In all cases a correct name was found.

We select first six names from the second column of `tax1`, by specifying this selection in brackets, and call `authors` to find their authorities:

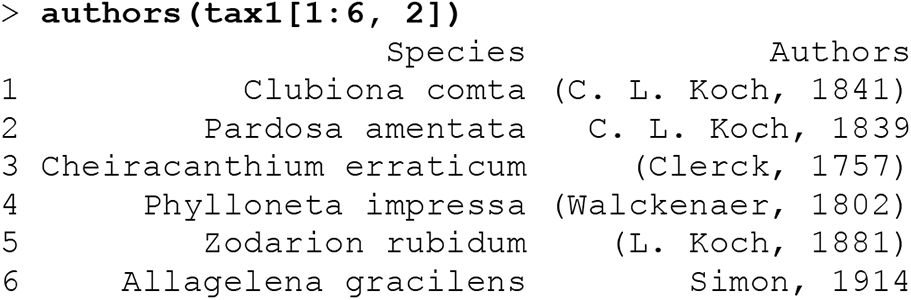

Authorities and years for all species in the list are given in the second column. Now we find the taxonomic classification for all these species using function `taxonomy`, without specifying any other arguments:

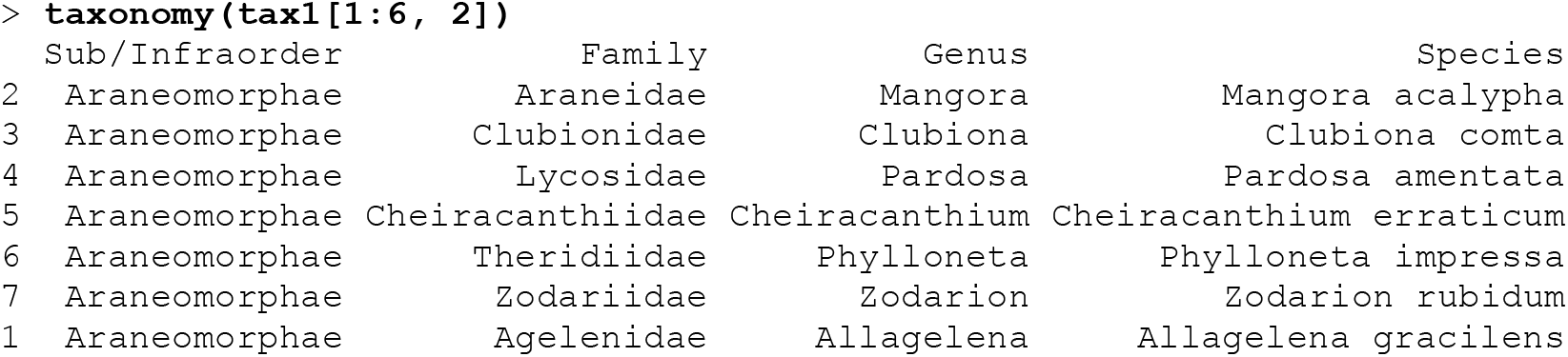

For each species name, the genus, family, and infraorder are shown. For the sake of completeness, we ask for LSID from WSC for all the species by calling `lsid`:

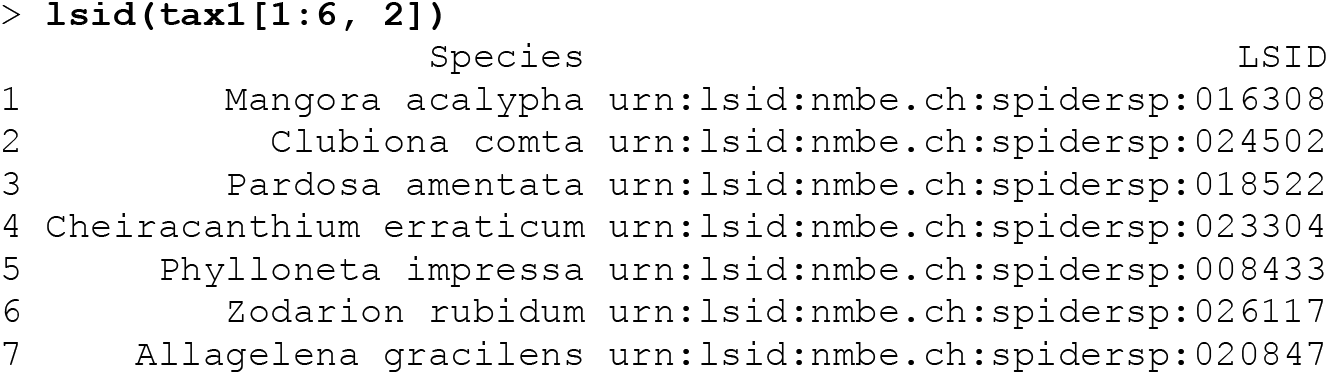

Eventually, we can get valid names, authorities, and classification in a single command by calling `taxonomy` with two arguments, `aut=T` and `check=T`. We will use the first seven species names from the original vector, tax. Results will be stored in an object, named e.g. `tax2`:

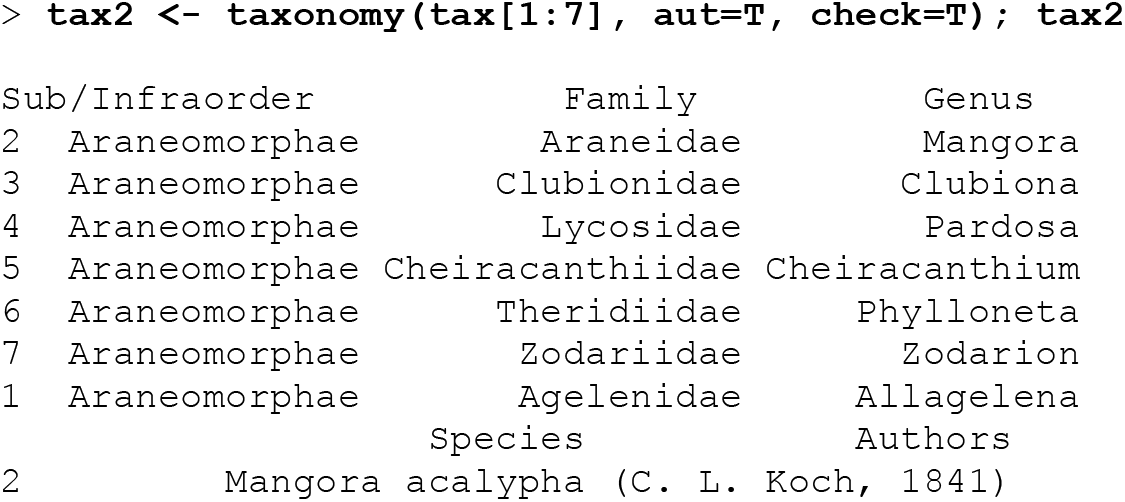
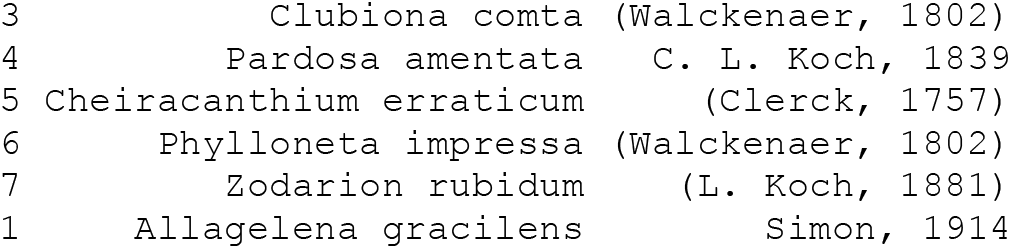

This output can be either marked and copied via clipboard into other software or exported to a file named e.g. table.txt on a disk (D) using the function `write.table`:

~~~
> **write**.**table(tax2, file=“d:\\table.txt“, sep=“\t”)**
~~~

### Example 2

In this example we show how to extract information on species and genus richness, how to gather data on a species distribution, and how to map them, which is often required for a taxonomy paper, for example.

If we need to know how many species there are in the family Palpimanidae and in the genus *Ero*, for example, we can use the function `length` and apply it on the function `species` with specified names:

~~~
> **length(species(“Palpimanidae”))**
[1] 158
**length(species(“Ero”))**
[1] 40
~~~

There are 158 species in the family Palpimanidae, and 40 species in the genus *Ero*. To obtain verbal description on the distribution of six species from the `tax1` we just type:

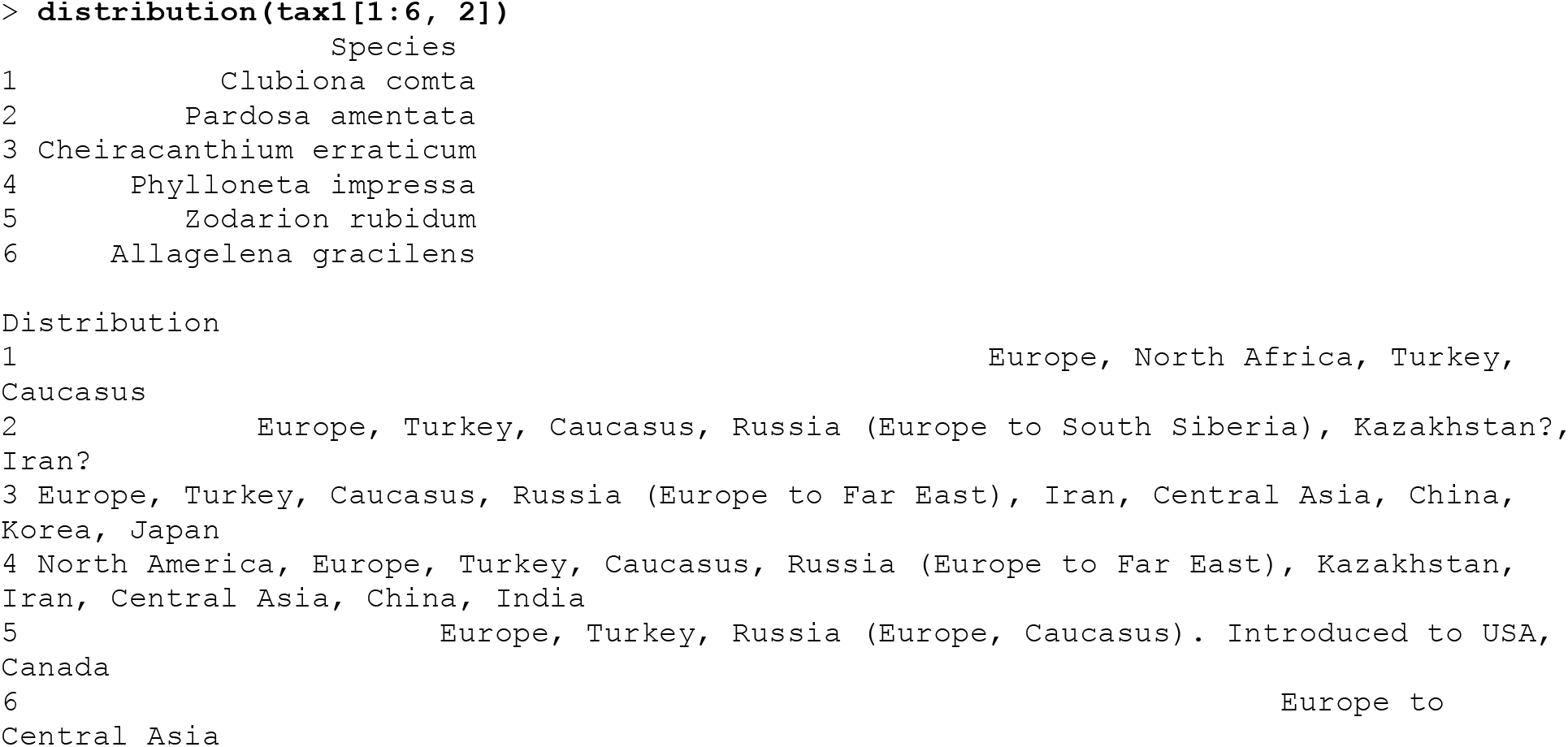

The verbal distribution information of several species is very long, thus it is returned on separate lines. Moreover, to obtain records for *Cheiracanthium erraticum*, which is listed among species in `tax1` at the 3^rd^ position in the second column, from WST and GBIF we call:

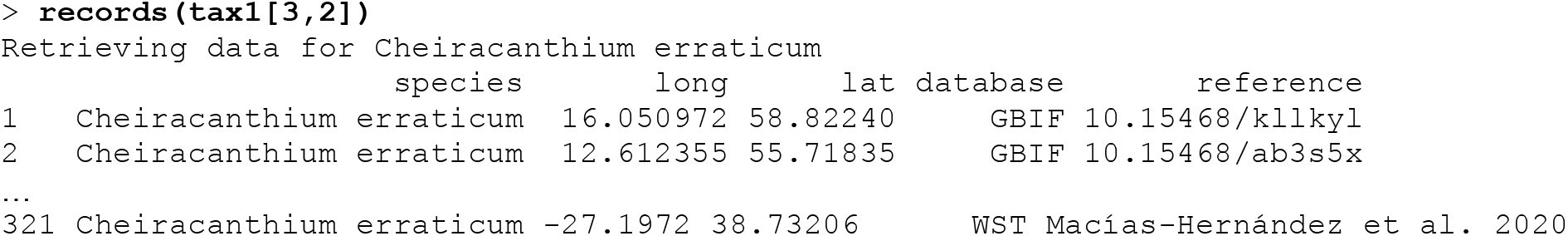

The list is very long, thus only the first two and the last row are displayed. Eventually, if we need to plot the species distribution, we use the function map. At first, we use a global world map of a species with a wide distribution, e.g. *Zodarion rubidum*, with countries of occurrence shaded, without records plotted as points (`records=F`).

~~~
> **map(“Zodarion rubidum”, records = F)**
~~~

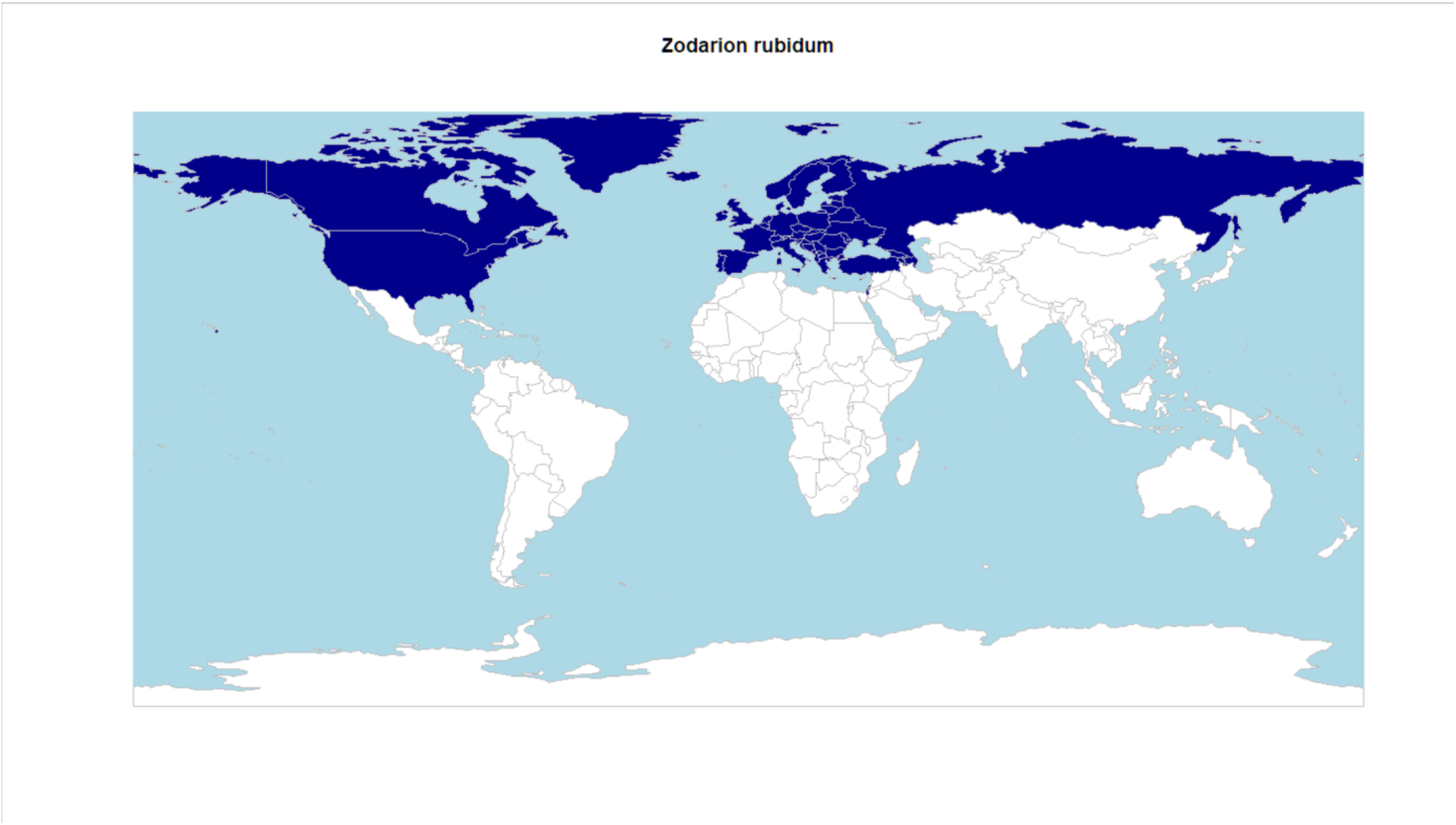

The user should be aware that the maps are only estimates of a species known distribution based on the WSC, with all the caveats associated with a rough textual description.

Secondly, we produce a map for a species with a small distribution, e.g., *Zodarion jozefienae*. For this purpose, we use the argument `zoom=T` and allow the actual records to be plotted as points. In order to get smooth country borders we use argument `hires=T`. Plotting might take longer.

~~~
> **<map(“Zodarion jozefienae”, hires = T, zoom = T)**
Retrieving data for Zodarion jozefienae
~~~

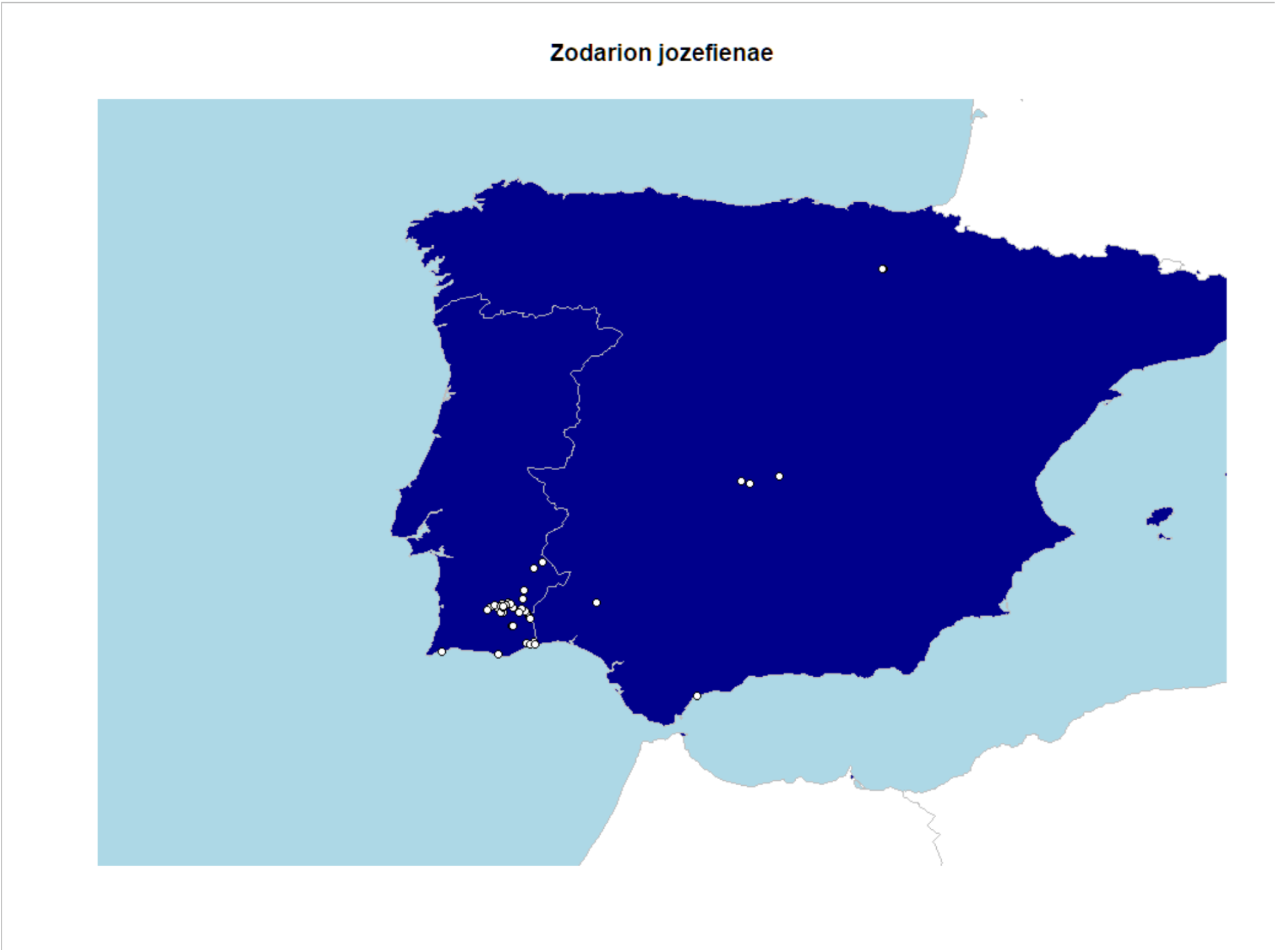

The function returned a map of the Iberian Peninsula with individual records.

### Example 3

Here we show how to prepare data on traits obtained from the WST for further statistical analysis. For example, we want to compare body length (abbreviation for this trait is “bole”) between males and females of *Zodarion jozefienae*. We download the trait data into an object named `tr1` by using the function traits with the species name as the first argument, and `trait = “bole”` as the second argument. Then we look at the content by calling names.

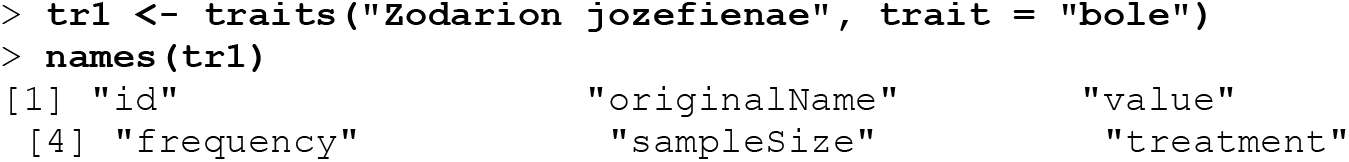
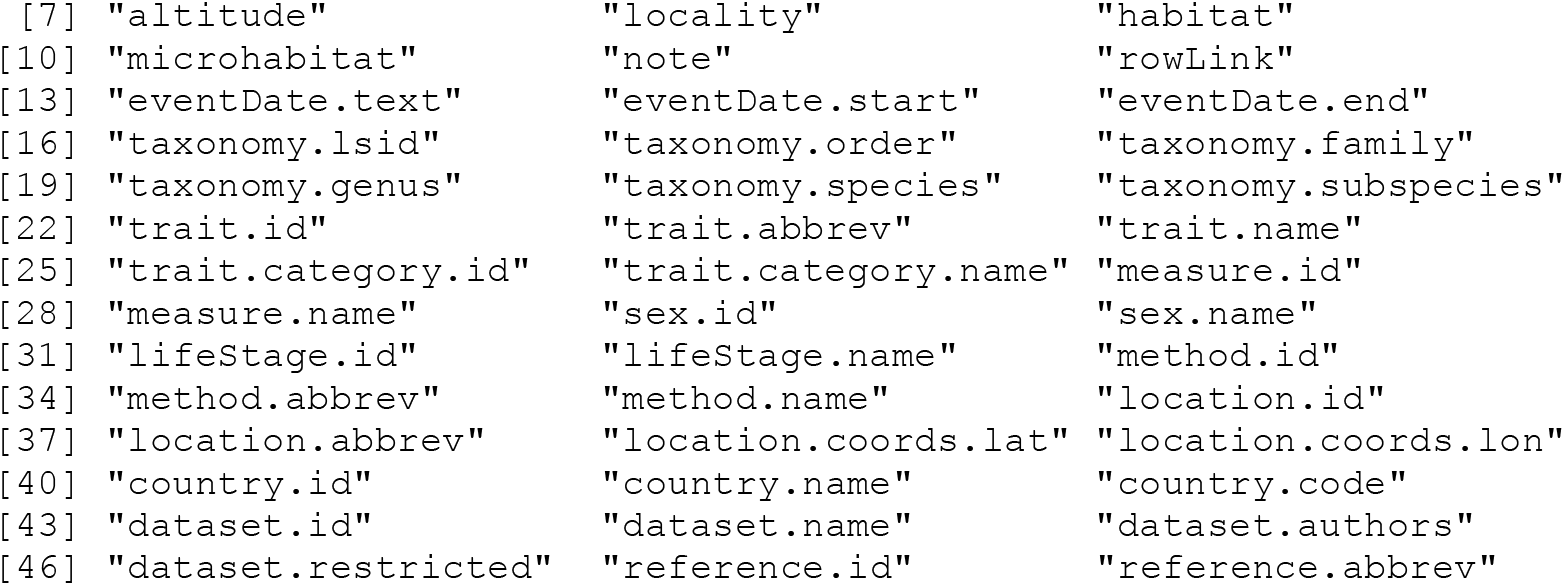

The data frame `tr1` has 48 columns. We are interested only in two of them, namely value (which contains the measurements) and sex.name (which contains the sex categories). For further analysis it is useful to extract those two variables from the `tr1` and it is essential to make the vector value numeric and the vector sex.name a factor. At last, we can produce a boxplot of the comparison.

~~~
> **body <-as**.**numeric(tr1$value)**
> **sex <-factor(tr1$sex**.**name)**
> **plot(body∼sex)**
~~~

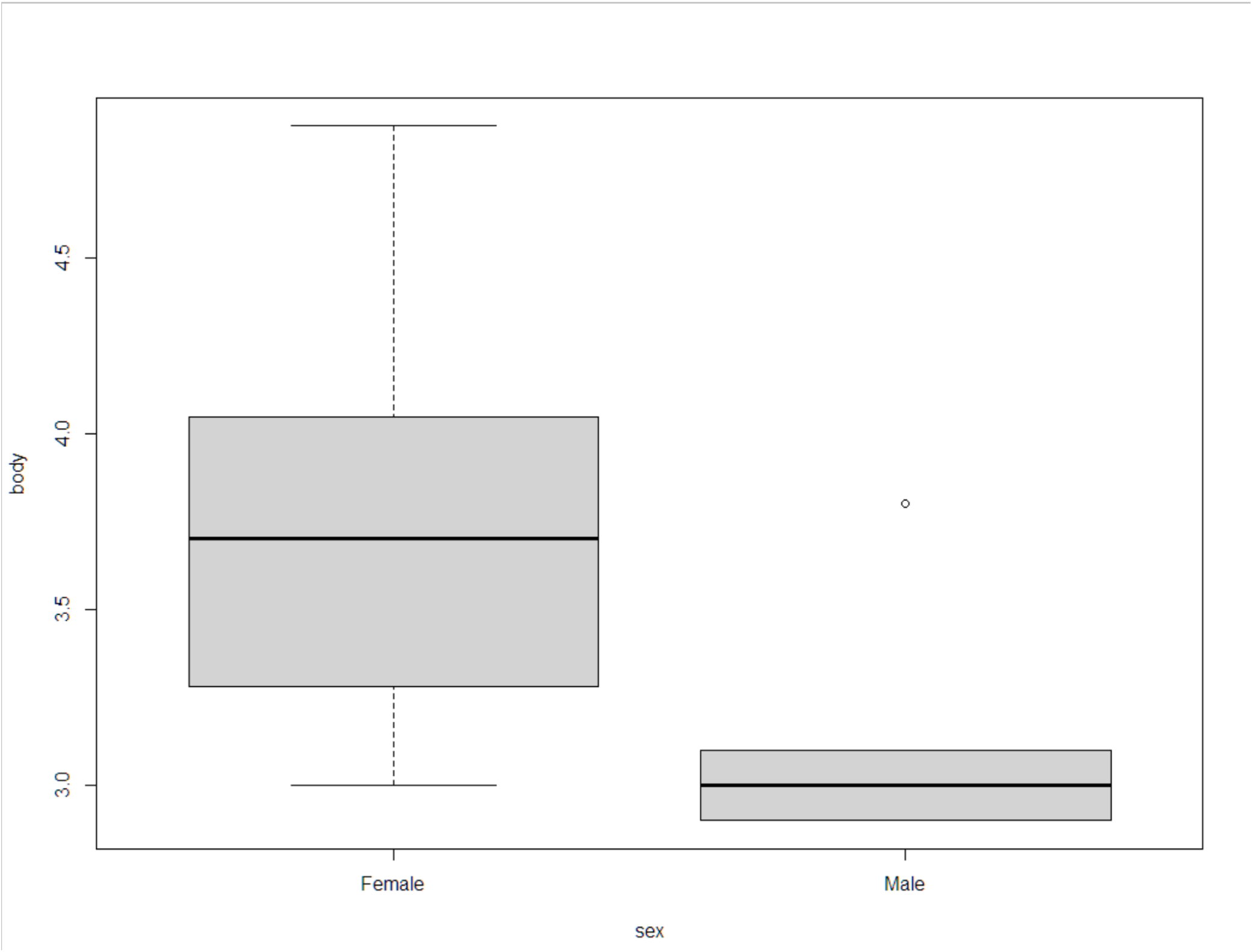

The boxplot shows larger body size in females than in males.

The two variables are ready to be used in statistical analyses, such as an ANOVA implemented in GLM.

